# Aspartate aminotransferase is required for *Salmonella* expansion in the inflamed gut via TCA anaplerosis

**DOI:** 10.64898/2026.02.24.707732

**Authors:** NG Shealy, M Baltagulov, HF Avalos, J Olivas, KM Jones, MX Byndloss

## Abstract

Aspartate represents an important proteogenic amino acid in all living organisms. Many microorganisms can produce aspartate through various biosynthetic processes, utilizing it for energy production and as a precursor for synthesizing other biomolecules, such as amino acids and nucleotides. The enteric pathogen *Salmonella* Typhimurium (*S*. Tm) has developed mechanisms to access aspartate as a nutrient source during expansion in the inflamed gut. However, how *S*. Tm deals with aspartate starvation during infection remains unknown. To address this knowledge gap, we interrogated *Salmonella*’s reliance on the bi-directional aspartate aminotransferase encoded by *aspC* for growth *in vitro* and during host colonization using murine models of *Salmonella* infection. AspC can interconvert aspartate and the TCA intermediate oxaloacetate and is hypothesized to support *S*. Tm cellular demands for aspartate during starvation or support refueling of the TCA cycle via oxaloacetate synthesis. Herein, we find that loss of *aspC* results in a gut-specific *S*. Tm colonization defect that increases with the course of infection. Importantly, *aspC* is dispensable for *S*. Tm systemic colonization in CBA/J mice. Additionally, we report that loss of *aspC* results in a significant growth defect during respiration of inflammation-derived electron acceptors *in vitro*. Interruption of oxidative TCA cycle progression via TCA enzyme deletion or supplementation with TCA intermediates (e.g., oxaloacetate) abrogates the defect observed in Δ*aspC S*. Tm *in vitro*. Thus, suggesting the requirement for AspC to catabolize aspartate and support the TCA cycle during respiration. Altogether, we report that AspC plays a critical role in *S*. Tm pathogenesis in a gut-specific manner during inflammation through supporting energy generation.

## INTRODUCTION

Despite being the primary site of digestion, the mammalian gut represents a region of nutrient scarcity for potential invading enteric pathogens such as *Salmonella enterica* serovar Typhimurium (*S*. Tm)^1–5^. The reason for this scarcity is the occupation of the major nutrient niches by the resident collection of commensal microbes inhabiting the intestines, known as the gut microbiota ^6–10^. The gut microbiota has been shown to actively reduce the availability of nutrients that potential invaders must compete for through direct catabolism and indirect support of host epithelial cell metabolism and nutrient absorption^11–13^. However, how bacterial pathogens adapt to overcome microbiota-protective mechanisms, such as nutrient sequestration, remains to be fully understood.

Recent work demonstrates that, in the context of complex microbiota, enteric bacterial pathogens such as *Citrobacter rodentium* and other pathogenic *Enterobacteriaceae* require the expression of amino acid biosynthesis pathways to colonize and expand^3^. Specifically, through a Tn-seq approach comparing germ-free and conventional mice, it has been demonstrated that mutants with insertions in amino acid biosynthesis pathways exhibit a significant fitness defect in conventional mice compared to germ-free mice. Thus, amino acids represent a critical nutrient for which microbial competition exists. Additionally, it is well-documented that bacteria are sensitive to amino acid starvation, which results in the activation of the stringent response. This response is commonly mediated by the SpoT/RelA system, which senses a reduction in amino acid-charged tRNAs, leading to reduced DNA replication, overall metabolism, and the upregulation of biosynthesis pathways. This body of work highlights the importance of the stringent response during gut colonization, as deleting these pathways in *S*. Tm results in a significant fitness defect and reduced virulence^14^. However, which amino acids are required for enteric pathogens in competition with complex microbiota is largely unknown.

Our group and others have previously reported that *S*. Tm requires aspartate for expansion in the inflamed gut through aspartate-dependent fumarate respiration via the aspartate ammonia-lyase, AspA ^15,16^. Specifically, deletion of *aspA* and the C4-dicarboxylic acid transporter encoded by *dcuA*, specific for aspartate import in *S*. Tm, results in a significant expansion defect during murine infection^15,16^. Thus, suggesting that aspartate plays a key role in the ability of *S*. Tm to colonize the intestinal lumen during gut inflammation. However, work by Popp *et al*. and Liu *et al*. demonstrates that aspartate biosynthesis also plays a significant role in *Salmonella* replication within macrophages and the maintenance of bacterial replication and cell size via the aspartate aminotransferase encoded by *aspC*^*17,18*^. Additionally, recent work by Vayena *et al*. through computational metabolic network mapping of *S*. Tm identified the potential for AspC to participate in the synthesis and breakdown of aspartate^19^. However, there is a knowledge gap in how aspartate prototrophy by *S*. Tm, more specifically the necessity of AspC, impacts the pathogen’s fitness during intestinal colonization, including during intestinal inflammation when the pathogen relies on anaerobic respiration to outcompete the resident microbiota.

Herein, we observe that deletion of the main aspartate aminotransferase AspC in *S*. Tm results in a significant gut-specific defect in pathogen expansion when comparing intragastric and intraperitoneal routes of infection in CBA/J mice. Based on our previous findings that *S*. Tm gastrointestinal infection results in an increased abundance of luminal aspartate^16^, we reasoned that the defect of Δ*aspC S*. Tm may not be the result of aspartate starvation but rather of defects in aspartate catabolism. AspC is known to be a bidirectional enzyme capable of degrading aspartate into the TCA intermediate oxaloacetate. Thus, we hypothesized that one possible reason for the expansion defect *in vivo* was the inability of Δ*aspC S*. Tm to fuel anaplerosis of the TCA cycle via oxaloacetate during anaerobic respiration. Specifically, during gut inflammation, the pathogen has increased access to both aspartate and inflammation-derived electron acceptors such as nitrate or tetrathionate, enabling *S*. Tm’s oxidative TCA cycle, using oxaloacetate for citrate synthesis. In line with this hypothesis, we find that *aspC*-deficiency results in significant growth and fitness defects only when *S*. Tm is respiring in an aspartate dose-dependent manner^20–22^. Using gene deletions in key TCA enzymes and supplementation of TCA intermediates, we confirm the role of AspC in supporting the central oxidative metabolism of *S*. Tm during anaerobic respiration. Altogether, this study deepens our understanding of the amino acid utilization by *S*. Tm, *in vivo*.

## RESULTS

Previous reports demonstrate that multi-organ site pathogens, such as *Staphylococcus aureus*, depend on *de novo* aspartate synthesis for survival in murine osteomyelitis models and other organ sites^23^. Transposon insertions in the aspartate aminotransferase of *S. aureus* result in decreased pathogen survival in the femoral bone in an osteomyelitis model of disease, as well as in the heart, kidneys, and liver of infected animals. Thus, suggesting the requirement for aspartate production by the pathogen for survival under the nutrient landscape of systemic organs. Another critical human pathogen is *S*. Tm, being one of the four leading causes of foodborne illnesses worldwide^24^. *S*. Tm is an enteric pathogen with disease centering in the cecum of mice^25,26^. However, *S*. Tm can also cause systemic disease by migrating to the liver, through biliary ducts, as well as the spleen, through hijacking of dendritic cells and spread through lymphatic vessels^27,28^. Additionally, the liver and spleen are known reservoirs for *Salmonella* replication in murine models of *S*. Tm-induced Typhoid-like disease and humans infected with *S*. Typhi and *S*. Paratyphi serovars^17,23,29^. Work by Nuccio *et al*. suggests that *aspC* is highly conserved amongst *Salmonella* serovars capable of causing systemic (Typhoid fever) or gastrointestinal disease (gastroenteritis), suggesting that aspartate biosynthesis may be required for *Salmonella* survival in systemic host sites during infection. However, there is a gap in knowledge regarding the dependence of aspartate aminotransferases broadly on the pathogens in different infection-relevant organ sites.

To address whether aspartate biosynthesis is required for *S*. Tm survival during systemic infection, we first constructed a *S*. Tm mutant lacking the major aspartate aminotransferase, encoded by *aspC*, and leveraged this strain to perform a competitive infection between wild-type (WT) *S*. Tm and the isogenic *aspC*-deficient strain (Δ*aspC*) in genetically resistant CBA/J mice. To do so, we infected CBA/J mice intraperitoneally with 10^4^ colony-forming units (CFU) of a 1:1 mixture of wild-type *S*. Tm and *S*. Tm Δ*aspC* (**Fig. 1A**), allowing the mice to carry the infection for three, four, or five-days before euthanasia. We chose these time points to understand if there were any temporal changes to the dependence of *aspC* by *S*. Tm before and after the onset of severe systemic disease. In line with our choice of time points, mice were maintained for five d.p.i. lost significantly more weight compared to those sacrificed at three d.p.i. (**Fig. 1B**). However, we observed no significant competitive advantage for WT *S*. Tm over the *S*. Tm Δ*aspC* mutant (**Fig. 1C**). While we observed that the overall *S*. Tm burden increased over time in all three organ sites (**Fig. 1D-F**), no differences in CFU recovered of each genotype were observed in any location following systemic infection (**Fig. 1D-F**). Together, these data suggest that, unlike *S. aureus, S*. Tm does not require the activity of AspC to survive in systemic sites during infection in the CBA/J mouse model.

**Figure 1:**
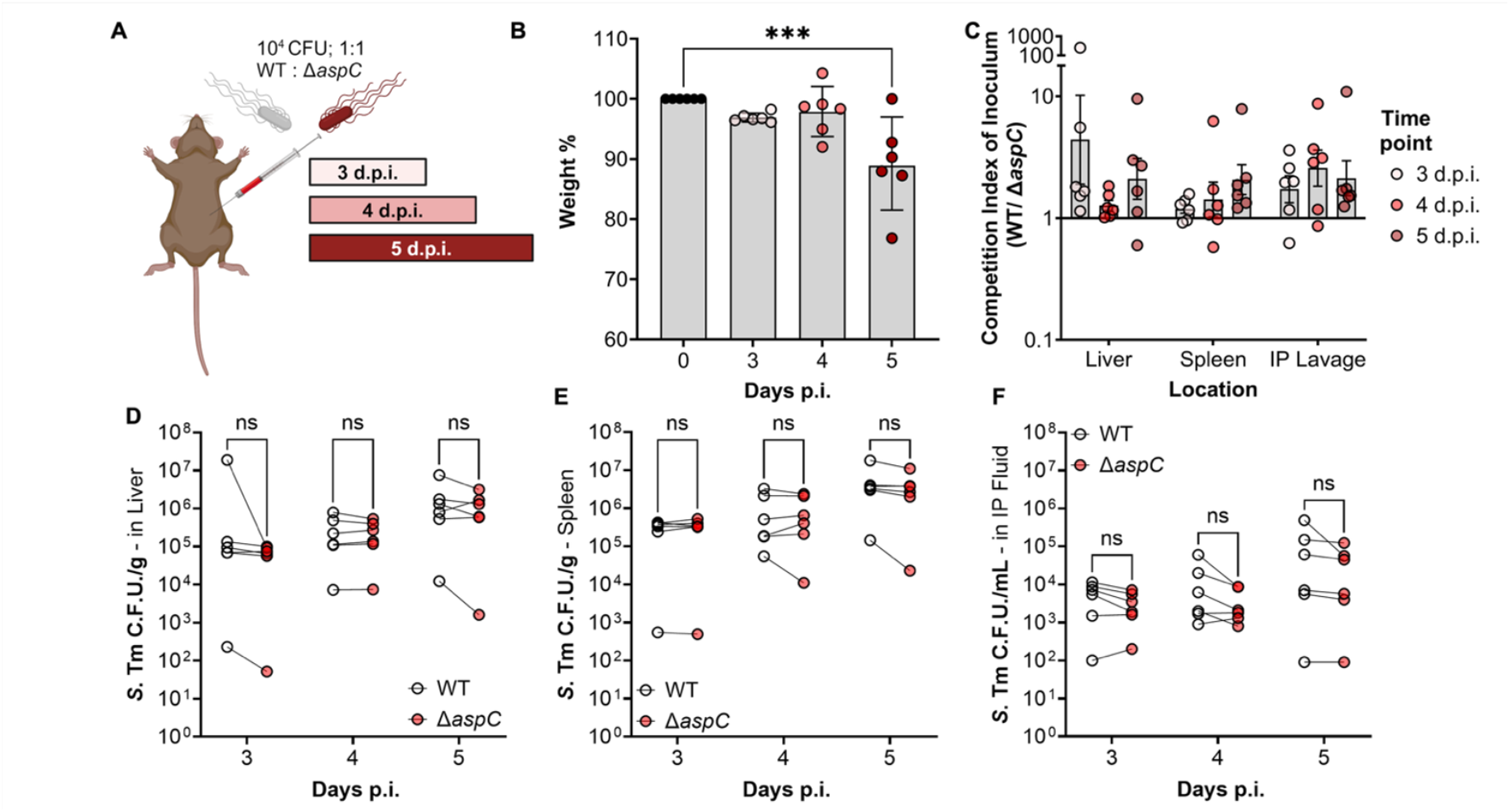
*S*. Tm does not require AspC for systemic survival following intraperitoneal injection. CBA/J mice were infected with a competitive 1:1 ratio of WT and *aspC*-deficient *S*. Tm IR715 (Δ*aspC*), via intraperitoneal injection at a dose of 10^4^ CFU. Mice were allowed to carry the pathogen for pre-determined time points, prior to human euthanasia and sample collection. (A) Infection Schematic. (B) Weight-loss percentage relative to inoculation weight in mice sacrificed three-, four-, and five-days post-infection. (C) Competitive index of inoculum at each time point based on the proportion of *S*. Tm in liver and spleen homogenate as well as intraperitoneal lavage fluid. (D) *S*. Tm burden of each genotype in the Liver. (E) *S*. Tm burden of each genotype in the CFU/g in spleen. (F) *S*. Tm burden of each genotype in the intraperitoneal fluid following lavage. N=6 per time point. Geometric mean with geometric SD. ***, p < 0.001 using multiple t-test.

### *S*. Typhimurium requires AspC during murine models of gastroenteritis

Before spreading to systemic sites, *S*. Tm colonizes and expands in the gastrointestinal tract^27^. We previously identified that *S*. Tm gains access to the amino acid aspartate during gut inflammation due to immune cell-mediated killing of the resident microbiota^16^. Secondly, we described how *S*. Tm leverages this newly available aspartate to generate fumarate via the aspartate ammonia-lyase AspA, supporting anaerobic respiration during intestinal inflammation^16^. However, how *S*. Typhimurium deals with aspartate limitation during initial colonization before gaining access to excess aspartate during intestinal inflammation remains to be fully elucidated. Given our observations of the dispensability of AspC during systemic infection, we explored the necessity of AspC for *S*. Tm host colonization during its native route of infection, the fecal-oral route. To do so, we orally gavaged CBA/J mice with 10^9^ CFU of a 1:1 mixture of WT *S*. Tm and isogenic *S*. Tm Δ*aspC*, allowing the animals to carry the infection for up to ten days (**Fig. 2A**).

**Figure 2:**
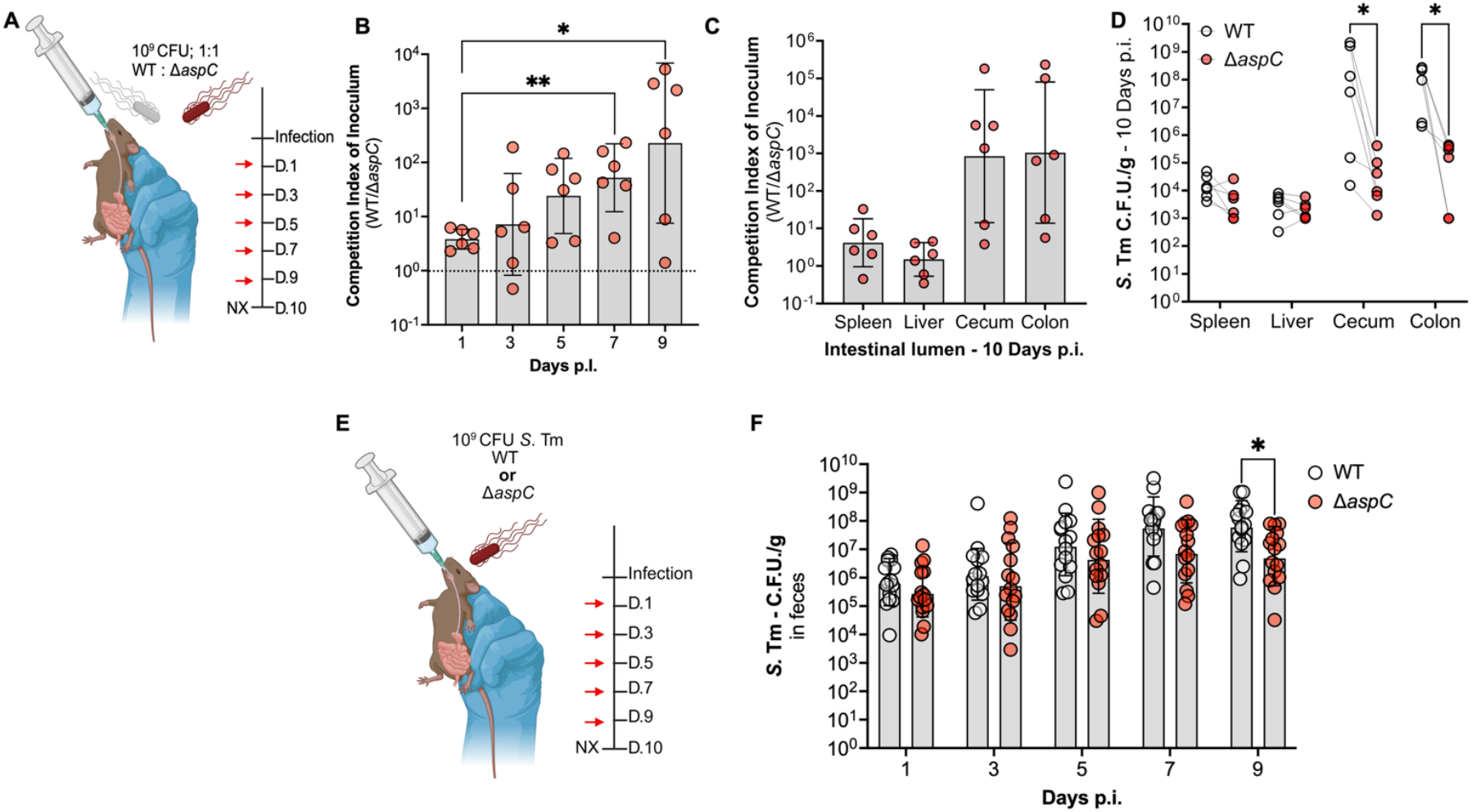
Deletion of *aspC* results in a significant expansion defect only in the cecum and colon, during murine model of gastroenteritis. To interrogate the necessity of AspC via the canonical fecal-oral route of infection, we gavaged CBA/J mice with a 1:1 competitive ratio of our WT *S*. Tm and isogenic Δ*aspC* mutant. (A) Schematic for competitive *in vivo* infection. (B) Competition index of inoculum over time in feces. (C-D) Competition index of inoculum ten d.p.i. from systemic and gut lumen samples of infected mice. To corroborate our observations, we repeated our infection with either of our strains in isolation. (E) Schematic for single infection. (F) *S*. Tm burden in feces over time in mice infected with either WT *S*. Tm or Δ*aspC* mutant. N=6 per condition. Geometric mean and Geometric SD. *, p < 0.05; **, p < 0.01 using Mann-Whitney (B), paired-end (E) or unpaired t-test (F).

Following initial intestinal colonization, we observed no defects of *S*. Tm Δ*aspC* compared to WT, as we recovered equal proportions of each genotype in feces one-day p.i. (**Fig. 2B**). However, as early as five days p.i., we observed a ten-fold colonization defect for *S*. Tm Δ*aspC* compared to WT in feces, which increased to ∼100-fold at nine days p.i. (**Fig. 2B**). Following sacrifice, we observed that the *S*. Tm Δ*aspC* defect was restricted to the intestinal lumen, with WT *S*. Tm outcompeting the isogenic Δ*aspC* mutant in the cecum and colon content (**Fig. 2C-D**). Additionally, AspC was not required for *S*.Tm survival in systemic sites such as the liver and spleen, as we recovered equal proportions of each *S*. Tm genotype, corroborating our IP infection observations (**Fig. 2C-D**).

To further interrogate the requirement for *asp*C during *S*. Tm gastrointestinal infection, we repeated our CBA/J mice infection model, now performing single infections. To do so, mice were orally infected with 10^9^ CFU of either WT *S*. Tm or isogenic Δ*aspC* mutant, and mice carried the infection for up to ten days (**Fig. 2E**). In line with our competitive infection results, we observed no defects in the Δ*aspC* mutant at one-day p.i. as we recovered similar CFU of WT and Δ*aspC* in feces. However, starting seven days p.i., we recovered significantly less Δ*aspC S*. Tm compared to WT in the feces of infected mice by nine days p.i. (**Fig. 2F**). Together, our findings suggest that *aspC* contributes to *S*. Tm intestinal colonization during gastroenteritis, specifically during expansion in the gut lumen in later stages of infection.

### Loss of T3SS does not significantly alter the dependence of *aspC* in *S*. Tm

Given our observation that the defect of Δ*aspC S*. Tm compared to the WT increased over time, in line with the onset of gastroenteritis around days seven to ten, we hypothesized this phenotype to be inflammation-dependent. To cause disease, *S*. Tm encodes two type III secretion systems, encoded by *Salmonella pathogenicity* islands 1 and 2 (T3SSs; SPI-1 & SPI-2), to invade the host epithelium and spread systemically, respectively^30,31^. Additionally, loss of both SPI-1 and SPI-2 renders *S*. Tm avirulent, unable to elicit gut inflammation^16^. To further interrogate the relationship between intestinal inflammation onset and the necessity of *aspC* for *S*. Tm intestinal colonization, we repeated our *in vivo* competitive infection in the CBA/J model of *S*. Tm-induced gastroenteritis, now using a *S*. Tm strain unable to induce intestinal inflammation through the genetic inactivation of T3SSs via deletions in *invA* and *spiB*^30,31^. Briefly, we orally gavaged CBA/J mice with 10^9^ CFU of a 1:1 mixture of Δ*invA*Δ*spiB S*. Tm and isogenic *S*. Tm Δ*aspC*, allowing the animals to carry the infection for up to ten days (**Fig. 3A**). Interestingly, compared to our wild-type background, genetic inactivation of the two T3SSs, increased the fitness defect of an *aspC-*deficient *S*. Tm, demonstrating a ten-fold defect significantly higher than that observed in the WT background one-day p.i. in the feces of mice (**Fig. 3B**). The competitive index between Δ*invA*Δ*spiB S*. Tm and isogenic Δ*invA*Δ*spiB*Δ*aspC* strains in fecal context remained steady throughout infection (between 10 to 100-fold) (**Fig. 3B**). Meanwhile, the competitive index between WT *S*. Tm and isogenic Δ*aspC* strain increased as the infection progressed, reaching 100-1,000-fold at day nine post-infection (**Fig. 3B**). While the competitive index between *S*. Tm WT and Δ*aspC* was significantly higher than the competitive index between Δ*invA*Δ*spiB* and its isogenic Δ*aspC* mutant in fecal samples at nine days post-infection (**Fig. 3B**), no significant differences are observed in the competitive index between the WT and Δ*invA*Δ*spiB* backgrounds in cecal (**Fig. 3C, E**) and colon (**Fig. 3D, F**) contents at 10 days p.i. In cecum and colon contents, both *S*. Tm WT and Δ*invA*Δ*spiB* strain significantly outcompeted their respective Δ*aspC*-deficient mutant to similar levels (**Fig. 3C-F**).

**Figure 3:**
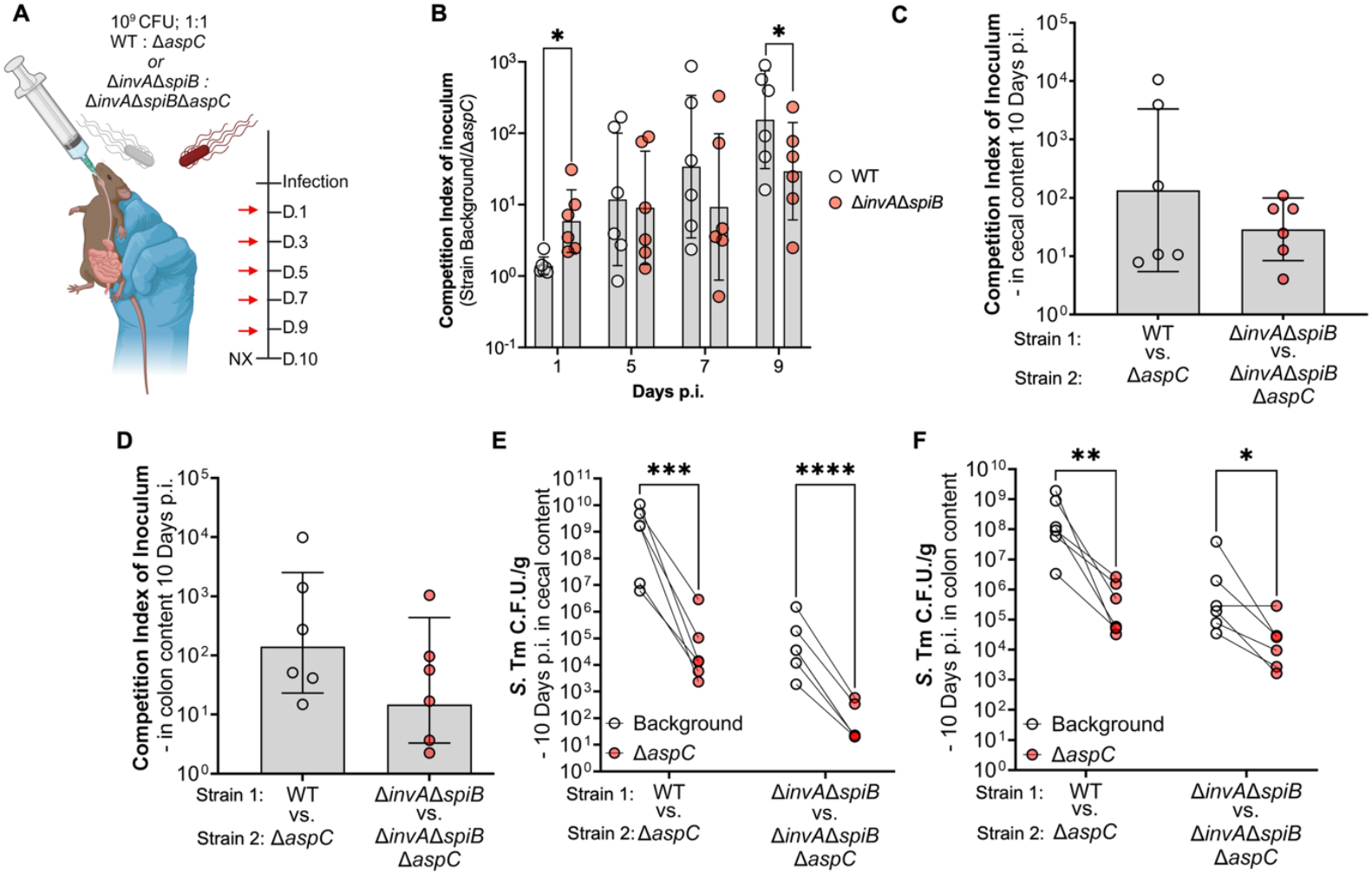
Genetic inactivation of T3SSs results in a colonization defect of Δ*aspC* but does not alter the overall expansion defect. We constructed isogenic *S*. Tm Δ*aspC* mutants in an avirulent background (Δ*invA*Δ*spiB*) and repeated the competitive infection experiment in CBA/J mice, comparing the avirulent strains to the *S*. Tm WT background. (A) Infection schematic of competitive infection. (B) Competitive index of inoculum in feces over the course of the infection. Day 3 samples were not collected. (C-D) Competitive index of inoculum (C) and colony forming unit (CFU) counts (D) of *S*. Tm strains in cecal content at 10 d.p.i.. (E-F) Competitive index of inoculum (E) and colony forming unit (CFU) counts (F) of *S*. Tm strains in colon content at 10 d.p.i. N=6. *, p < 0.05; **, p < 0.01; ***, p < 0.001; ****, p < 0.0001 using unpaired- (B) or paired t-test (E, F).

Given that the *S*. Tm Δ*aspC* colonization defect increases over time (**Fig. 3B**), which correlates with the onset of intestinal inflammation, we hypothesized that AspC may be more critical during anaerobic respiration, which *S*. Tm relies upon for expansion in the gut lumen during intestinal disease. To explore this hypothesis, we cultured WT *S*. Tm and our *aspC*-deficient mutant anaerobically *in vitro* under conditions that model the healthy and inflamed gut. Specifically, we grew WT *S*. Tm and Δ*aspC* in LB media or NCE minimal media supplemented with glucose as a carbon source or in NCE minimal media supplemented with glycerol as a carbon source in the presence of aspartate and inflammation-derived electron acceptors such as nitrate, as we have previously performed ^16^. Neither strain demonstrated an overall growth defect in the presence of LB nor NCE supplemented with glucose as a carbon source, which are conditions mimicking the healthy intestine (**Fig. 4B; S1A**). However, we observed an aspartate-dependent growth defect in the *S*. Tm Δ*aspC* strain upon culture in NCE supplemented with nitrate, using glycerol as a carbon source, which mimics the inflamed gut (**Fig. 4C, D**), and the anaerobic *S*. Tm growth under glycerol and nitrate conditions was restored when the *aspC* gene was reintroduced into the *S*. Tm Δ*aspC* genome (**Fig. S1B-C**). Specifically, *S*. Tm Δ*aspC* demonstrated growth defects when growing under aspartate-limiting conditions (0 or 0.1mM aspartate). However, the *S*. Tm Δ*aspC* growth defect was ameliorated through the addition of 10mM aspartate (**Fig. 4D**). We have detected similar changes in magnitude in the abundance of aspartate in the gut lumen during *S*. Tm infection, increasing from 1µM/g up to 10µM/g of feces ^16^. Similar results were obtained under conditions supplemented with either tetrathionate (**Fig. 4E**) or under hypoxia (**Fig. S1D**) as electron acceptors. To corroborate our observations from growth assays, we performed an *in vitro* competitive infection comparing the fitness of *S*. Tm WT versus the isogenic Δ*aspC* mutant under anaerobic respiration conditions. In line with our growth assay, we observed that *S*. Tm Δ*aspC* is significantly outcompeted by the WT strain in aspartate-limited conditions in which the bacterium was anaerobically respiring inflammation-derived electron acceptors (**Fig. 4F, G**). The fitness defect of *S*. Tm Δ*aspC* during anaerobic respiration could be abrogated by adding 10 mM aspartate. Previous work in our group and others demonstrated that exposure of *S*. Tm to aspartate significantly increases the expression of aspartate catabolizing enzyme AspA, dependent on the two-component system DcuRS^15,16^. To account for potential differences in the expression of *aspC* in the conditions we tested, we assayed *aspC* expression in *S*. Tm grown anaerobically with and without nitrate or 10 mM aspartate (**Fig. 4H**). No significant differences in *aspC* expression were detected between the conditions tested (**Fig. 4H**). Together, these data demonstrate that a loss of *aspC* significantly hinders the ability of *S*. Tm to anaerobically respire *in vitro*.

**Figure 4:**
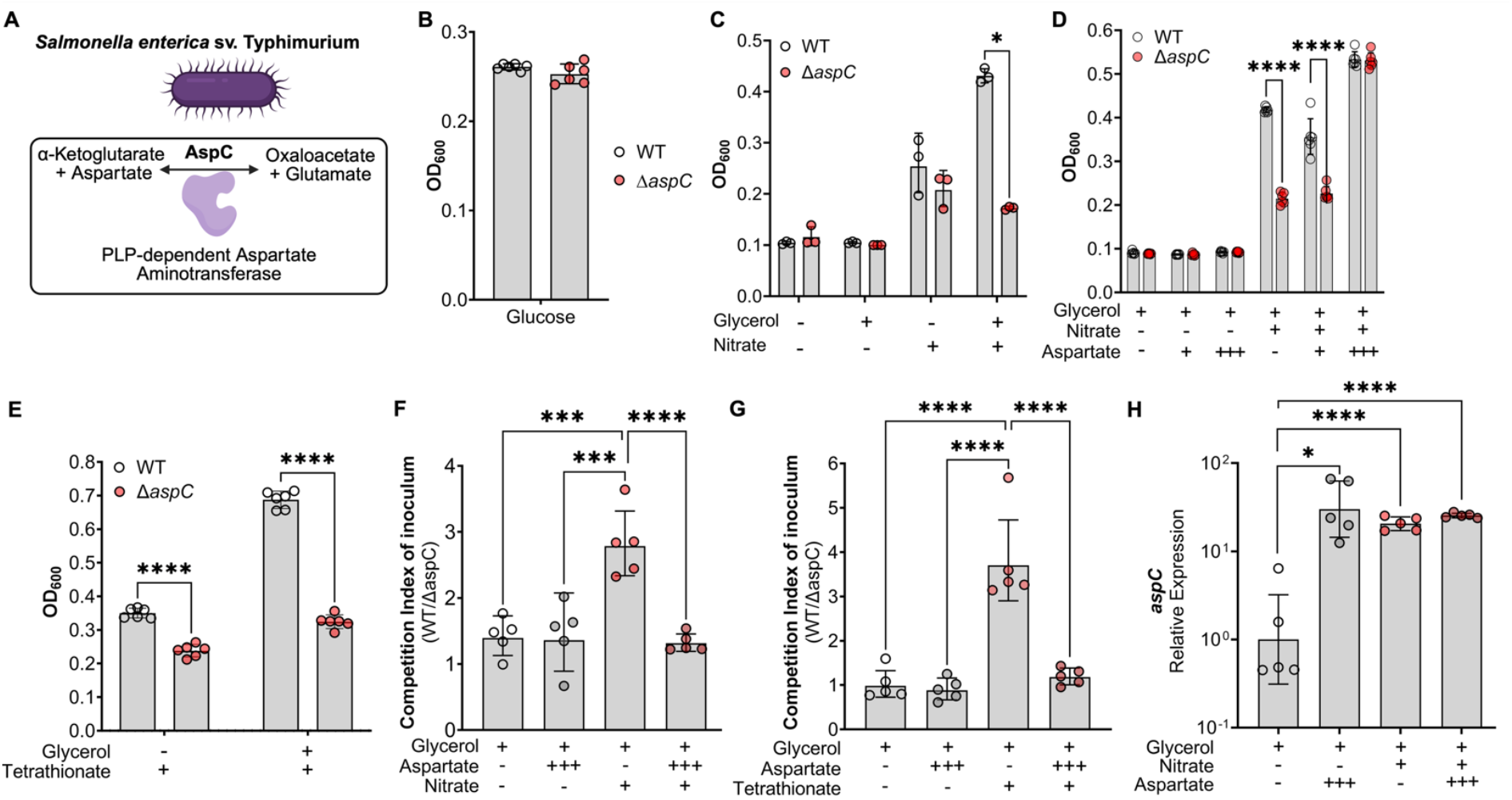
AspC is required for *S*. Tm growth under aspartate-limited conditions, confers a significant fitness cost during anaerobic respiration. (A) Schematic of the enzymatic role of AspC in *S*. Tm. (B) Anaerobic growth of WT and *aspC*-deficient *S*. Tm in NCE minimal media supplemented with 5 mM glucose at 18 hours. (C) Anaerobic growth of WT and *aspC*-deficient *S*. Tm (Δ*aspC*) in NCE minimal media supplemented with 40mM Glycerol with or without 40mM Nitrate at 18 hours. (D) Anaerobic growth of WT and *aspC-*deficient *S*. Tm (Δ*aspC*) in NCE minimal media supplemented with (+) 0.1mM or (+++) 10mM Aspartate with Glycerol and Nitrate at 18 hours. (E) *In vitro* competition of WT vs *aspC* in NCE supplemented with 40mM Tetrathionate, 10mM Aspartate, and 40mM Glycerol at 18 hours. (F) *In vitro* competition of WT vs Δ*aspC* in NCE supplemented with 40mM Nitrate, 10mM Aspartate, and 40mM Glycerol at 18 hours. (G) Relative expression of *aspC* in WT *S*. Tm cultured anaerobically for 4 hours in media supplemented with 10mM Aspartate, 40mM Glycerol, and 40mM Nitrate. N=3-10 per condition, at least 2 independent experiments for each, Geometric mean with geometric SD. *, p < 0.05; ****,p < 0.0001 using multiple t-test (C-E) or one-way ANOVA (F-H).

### Loss of AspC results in altered biosynthetic processes in *S*. Tm during nitrate respiration

To further explore the requirement of AspC during nitrate respiration, we compared the gene expression profile of *S*. Tm WT to Δ*aspC* cultured in minimal media. Given the critical role of aspartate in the central metabolism of *S*. Tm, we aimed to interrogate how aspartate deficiency influenced biosynthesis processes that rely upon aspartate as a precursor. Specifically, work done in *E. coli* and *S*. Tm demonstrates that these closely related bacteria can use aspartate as a precursor for the synthesis of many biological molecules, such as amino acids and nucleotides (**Fig. S2A**). We were specifically interested in assessing differences in the biosynthetic pathways in conditions that model the inflamed gut, as these are the conditions in which we identify an *aspC-*dependent growth and fitness defect. To do so, we cultured either WT *S*. Tm or the isogenic Δ*aspC strain* in NCE minimal media supplemented with 40 mM glycerol and 40 mM nitrate to evaluate differences in gene expression between genotypes. First, we were able to confirm our genetic inactivation, as Δ*aspC* demonstrated a significant reduction in the expression of *aspC* (**Fig. S2A-B)**. Additionally, we found that Δ*aspC* significantly upregulates the aspartate importer encoded by *dcuA* but not the aspartate ammonia-lyase encoded by *aspA* (**Fig. S2C-D**). Lastly, we report that loss of *aspC* in the presence of nitrate results in a significant increase in the expression of genes associated with pyrimidine synthesis, such as *pyrB* and the amino acid lysine, *lysC* (**Fig. S2E-F)**. No differences were observed between WT and Δ*aspC* in the expression of genes associated with methionine synthesis, specifically *metL* (**Fig. S2G**). Altogether, these data suggest that genetic inactivation of *aspC* alters the cellular demands of the bacterium, specifically increasing expression of genes associated with aspartate uptake and synthesis of aspartate-dependent biomolecules (e.g., pyrimidines and lysine). However, our gene expression analysis indicates that aspartate-dependent biosynthetic pathways remain functional in the absence of AspC.

### AspC supports the central metabolism of *S*. Tm during anaerobic respiration

Based on the observation that deletions in *aspC* significantly reduce *S*. Tm fitness during anaerobic respiration (**Fig. 4**) as well as the limited impact of *aspC* deletion on aspartate-dependent biosynthesis pathways in *S*. Tm *in vitro* under anaerobic respiration conditions (**Fig. S2E-G**), we hypothesized that AspC’s role may not be related to aspartate production. Instead, AspC may be required for *S*. Tm conversion of aspartate into oxaloacetate (**Fig. 5A**). To explore the role of AspC in supporting *S*. Tm anaerobic respiration through oxaloacetate production, we used a combination of *in vitro* supplementation using TCA intermediates and genetic inactivation studies. First, we repeated our *in vitro* competitions between WT *S*. Tm and isogenic Δ*aspC* strain in NCE media supplemented with 0.5% porcine mucin. We chose to perform our competitions using NCE + 0.5% porcine gastric mucin to enable us to adequately model the conditions of the inflamed gut, utilizing a physiologically relevant nutrient source (e.g., mucin), which allows *S*. Tm to ferment or respire available carbon sources in the media. Additionally, we supplemented the NCE + 0.5% porcine mucin media with either 10 mM fumarate, 10 mM succinate, or 10 mM oxaloacetate in the presence or absence of nitrate. We specifically chose these TCA intermediates as they are used during both the branched chain TCA and oxidative TCA cycles (**Fig. 5A**). Like our previous observations in NCE minimal media supplemented with glycerol, WT *S*. Tm significantly outcompeted Δ*aspC* only in mucin media supplemented with inflammation-derived electron acceptors such as nitrate (**Fig. 5B-D**). If the role of AspC is to support *S*. Tm’s ability to run the TCA cycle via oxaloacetate production, supplementation of the media with other TCA intermediates should abrogate the *S*. Tm Δ*aspC* growth defect under anaerobic respiration conditions. To test this hypothesis, we performed an *in vitro* competition in NCE + 0.5% porcine mucin media supplemented with 10 mM fumarate, succinate, or oxaloacetate in the presence or absence of nitrate. Notably, the addition of either of these TCA cycle intermediates to mucin media did not result in a fitness defect of the Δ*aspC* mutant compared to the WT strain in the absence of nitrate (**Fig. 5B-D**). However, under anaerobic respiration conditions (e.g., mucin media + nitrate), the addition of fumarate (**Fig. 5B**), succinate (**Fig. 5C**), or oxaloacetate (**Fig. 5D**) resulted in a significant reduction in the competitive advantage of the WT *S*. Tm over the Δ*aspC* mutant when compared to mucin media supplemented with nitrate alone (**Fig. 5B-D**). Altogether, these data suggest that loss of *aspC* results in a deficiency in TCA intermediates, namely oxaloacetate, which may lead to a reduction in energy production potential during anaerobic respiration.

**Figure 5:**
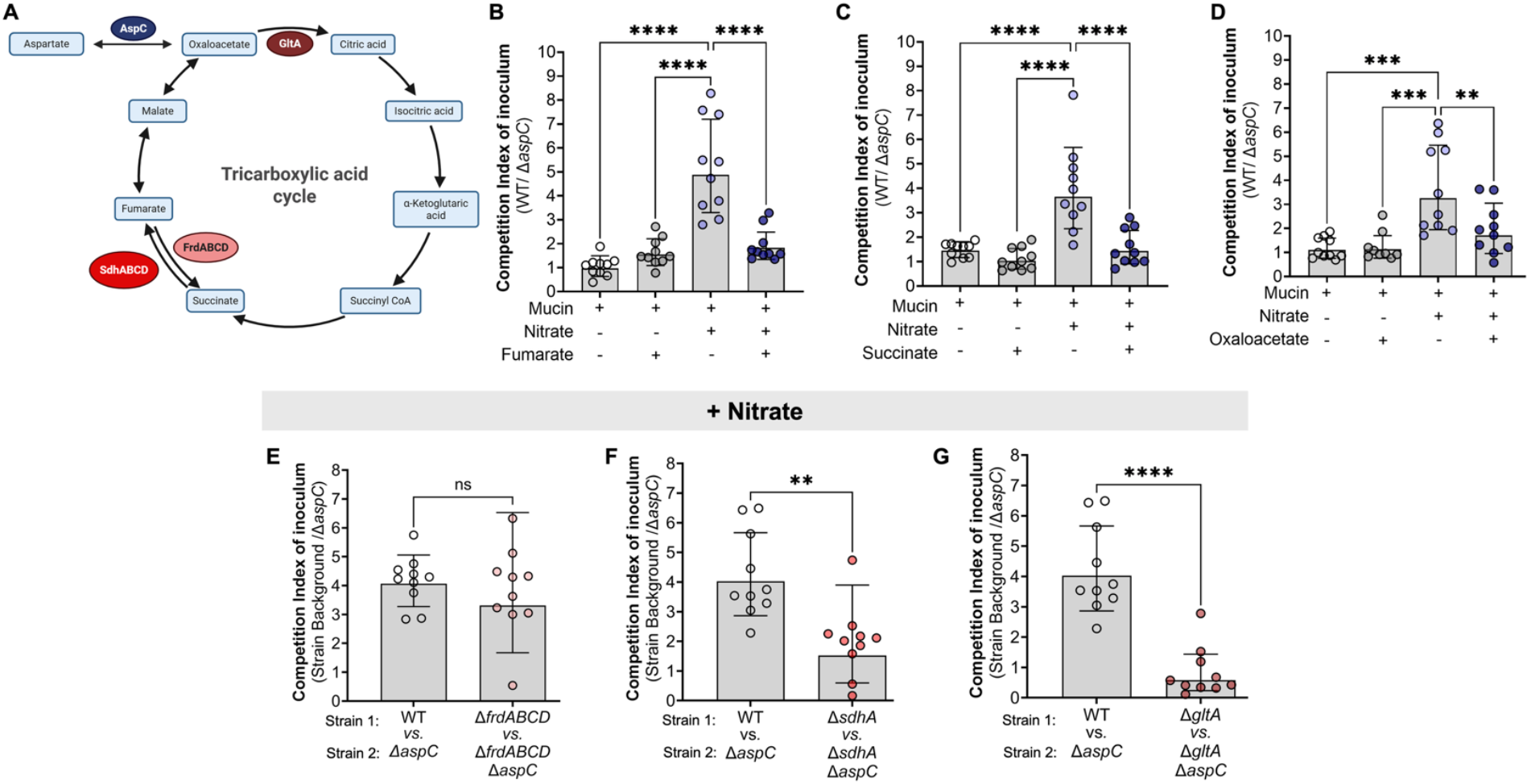
AspC is required for anaplerosis of the TCA by *S*. Tm during anaerobic respiration, *in vitro*. (A) Tricarboxylic acid cycle. (B-D) To confirm the role of AspC during nitrate respiration, we performed competitive infections *in vitro* under anaerobic conditions in the presence or absence of TCA intermediates. (B-D) Competitive index of inoculum at 18 hours between WT and Δ*aspC S*. Tm in the presence or absence of nitrate supplemented with (B) Fumarate (C) Succinate, and (D) Oxaloacetate. (E-F) We repeated our *in vitro* competitive infections between WT *S*. Tm and Δ*aspC* mutant during nitrate respiration using *S*. Tm backgrounds lacking the enzymes responsible for the utilization of: (E) Fumarate via fumarate reductase (Δ*frdABCD*); (F) succinate via succinate dehydrogenase subunit A (Δ*sdhA*); (G) oxaloacetate via citrate synthase (Δ*gltA*). N= 10 per condition. *, p < 0.05; **, p < 0.01; ***, p < 0.001; ****,p < 0.0001 using One-way ANOVA (B-D) or unpaired t-test (E-G).

Before the onset of inflammation, *S*. Tm has been shown to rely upon the branched-chain TCA, which is supported by fumarate respiration via the fumarate reductase FrdABCD^15^. However, once intestinal inflammation is established and electron acceptors become available, *S*. Tm relies on the canonical oxidative TCA cycle through the coordinated expression of TCA enzymes such as *sdhABCD* and *gltA* ^32^. To further interrogate the relationship between the progression of either the reductive branched-chain or oxidative TCA cycle and the defect of the Δ*aspC S*. Tm mutant during anaerobic respiration, we constructed mutants lacking the citrate synthase, *gltA*, the first component of the succinate dehydrogenase, *sdhA*, and the fumarate reductase, *frdABCD*. These genes represent significant steps of the branched chain reductive (FrdABCD) and oxidative (GltA & SdhA) TCA cycles in *S*. Tm (**Fig. 5A**). To understand the role of AspC in the TCA enzyme-deficient strain backgrounds, we constructed *S*. Tm isogenic mutants lacking *aspC*, Δ*gltA*Δ*aspC*, Δ*sdhA*Δ*aspC*, and Δ*frdABCD*Δ*aspC*, respectively. We hypothesized, based on our supplementation studies, that interruption of the TCA cycle progression via deletions in *gltA* and *sdhA* but not *frdABCD* would abrogate the defect of *S*. Tm Δ*aspC* during nitrate respiration. To test this hypothesis, we performed an *in vitro* competition in 0.5% porcine mucin media in the presence or absence of nitrate, specifically competing *S*. Tm WT, Δ*frdABCD*, Δ*sdhA*, Δ*gltA* with their respective isogenic *aspC* mutants.

First, to assess the requirement for *aspC* during the reductive branched-chain TCA cycle in *S*. Tm, we compared the competition index of our mutant Δ*frdABCD*: Δ*frdABCD*Δ*aspC* with that of WT:Δ*aspC* in minimal mucin media without nitrate. Interestingly, we observed a significantly increased fitness defect of Δ*frdABCD* Δ*aspC* compared to the Δ*frdABCD* isogenic control (CI∼10) (**Fig. S3A**) over that of WT vs. Δ*aspC* (CI∼1). In line with the literature that *frdABCD* is significantly downregulated during exposure to nitrate, we observed no significant difference in the fitness defect of an Δ*aspC* mutant during nitrate respiration in the *S*. Tm background lacking *frdABCD* (**Fig. 5E**). Together, these data suggest that AspC supports the branched chain TCA cycle during which fumarate respiration occurs and that FrdABCD is not required for the AspC-mediated effect on the *S*. Tm oxidative TCA cycle during nitrate respiration.

To determine the requirement for *sdhA* and *gltA* to the AspC-mediated support of *S*. Tm growth during the branched chain TCA cycle, we compared the competition index of our mutants Δ*sdhA*: Δ*sdhA*Δ*aspC* or Δ*gltA*: Δ*gltA*Δ*aspC* to WT: Δ*aspC* in 0.5% porcine mucin media without nitrate. Loss of *aspC* conferred no fitness defect in *S*. Tm strains lacking *sdhA* and *gltA* (**Fig. S3B**). Conversely, loss of *sdhA* or *gltA* significantly abrogated the fitness defect of a *S*. Tm Δ*aspC* mutant when compared to the isogenic WT background when strains were grown in 5% porcine mucin media supplemented with nitrate, a condition that enables *S*. Tm to run the oxidative TCA cycle (**Fig. 5F-G**). Together, our results using a genetic approach relying on the deletion of genes in *S*. Tm involved in the TCA cycle complement the results from the TCA cycle intermediate supplementation experiments (**Fig. 5B-D**) and pinpoint the requirement for *aspC* to support the progression of the oxidative but not reductive branched-chain TCA cycle in *S*. Tm.

## DISCUSSION

Herein, this study establishes a role for the aspartate aminotransferase, AspC, in the expansion of *Salmonella* Typhimurium in the mammalian gut. Using intraperitoneal and intragastric infection routes, we report that in the CBA/J mouse model of *S*. Tm infection, AspC is required only during intestinal expansion and is dispensable at systemic sites such as the liver and spleen. The role of AspC in *S*. Tm-induced systemic disease in susceptible mouse models (e.g., C57Bl/6) remains to be determined. Additionally, we find that the necessity of AspC for *S*. Tm intestinal survival is potentially independent of the main virulence program of *S*. Tm, the two T3SSs. However, we observe a loss of the ability to invade, and intracellular replication significantly increases the defect in an isogenic *aspC-*deficient strain at 1 d.p.i. Thus, our data suggest that *S*. Tm may subvert the initial response to aspartate starvation in the healthy gut by invading the gut epithelium. This hypothesis is in line with work by Popp *et al*., which demonstrates that AspC is not required for invasion or intracellular replication in HeLa cells^18^. Granted, there may be cell-type-specific differences in nutrient availability between uterine and intestinal epithelial cells, and future work should aim to understand the nutrient requirements between different infection-relevant organ sites.

Our study indicates that a loss of *aspC* results in altered metabolic demands for the pathogen, as determined by qPCR, *in vitro* supplementation, and genetic inactivation of key TCA enzymes. We observe that loss of AspC results in increased expression of genes encoding enzymes associated with the import and utilization of aspartate for the synthesis of amino acids and nucleotides. However, these alterations have a limited induction, resulting in less than a 2-fold change, suggesting tight control of these central metabolic processes. Alternatively, we find that *S*. Tm Δ*aspC* exhibits a significant growth defect under *in vitro* conditions that promote the reductive branched-chain and oxidative TCA cycles. Thus, this suggests one possible explanation for the defect in *S*. Tm Δ*aspC* in gut expansion: the inability of this strain to exploit the disruption in the gut metabolic landscape during gastroenteritis.

Lastly, while we do not observe major differences in the competitive fitness of *S*. Tm Δ*aspC* in a background lacking *frdABCD* under *in vitro* conditions supporting anaerobic respiration, we cannot rule out the possibility of the fumarate reductase FrdABCD aiding in driving the oxidative TCA forward, as its expression is induced in *S*. Tm under *in vitro* anaerobic growth conditions characterized by the presence of excess aspartate and nitrate which resemble the growth conditions encountered by the pathogen in the inflamed gut^16^. Future work should aim to untangle the relationship between FrdABCD and SdhABCD in their roles in supporting the pathogenesis and expansion of *S*. Tm in the mammalian gut.

Altogether, our findings expand our understanding of *S*. Tm aspartate metabolism and its influence on survival in different organ sites. Future studies on bacterial metabolism are poised to elucidate previously unknown aspects of pathogenic bacterial survival and inform therapeutic strategies targeting these essential metabolic pathways.

## MATERIALS & METHODS

### Resources availability

#### Lead contact

Further information and requests for resources and reagents should be directed to and will be fulfilled by the lead contact, Mariana X. Byndloss.

#### Materials availability

All unique reagents generated in this study are available from the lead contact without restriction.

#### Data and code availability

Data reported in this paper will be shared upon request by the lead contact upon request. This paper does not report original code. Any additional information required to reanalyze the data reported in this paper is available from the lead contact upon request.

### In vitro

#### Bacterial culture

*S*. Typhimurium IR715 strains were routinely grown aerobically at 37°C in LB broth (10 g/L tryptone, 5 g/L yeast extract, 10 g/L sodium chloride) or on LB agar plates (10 g/L tryptone, 5 g/L yeast extract, 10 g/L sodium chloride, 15 g/L agar). When appropriate, agar plates and media were supplemented with 30 μg/mL chloramphenicol (Cm), 100 μg/mL carbenicillin (Carb), 100 μg/mL kanamycin (Km), or 50 μg/mL nalidixic acid (Nal). For growth under anaerobic conditions, No-carbon-E (NCE) minimal medium (28 mM KH_2_PO_4_, 28 mM K_2_HPO_4_⋅3H_2_O, and 16 mM NaNH_4_HPO_4_⋅4H_2_O) supplemented with 1 mM MgSO_4_, 0.1% Casamino acids, and 1% Vitamin and Trace mineral solutions (American Type Culture Collection) was used. As a sole carbon source, glucose (5 mM) or glycerol (40 mM) was added to the bacterial culture in addition to L-aspartate (0.1-10 mM). When stated otherwise, bacteria were cultured in Mucin minimal media, NCE minimal media (28 mM KH_2_PO_4_, 28 mM K_2_HPO_4_⋅3H_2_O, and 16 mM NaNH_4_HPO_4_⋅4H_2_O) supplemented with 0.5% Porcine Gastric Mucin (Type II) (ThermoFischer) supplemented with 40mM sodium nitrate, 10mM sodium fumarate, succinic acid, or oxaloacetic acid.

#### Mutant generation

Mutant construction was performed via lambda red recombination as previously described ^34^. In brief, primers were designed to include 40bps homologous regions upstream and downstream of the gene of interest and used to amplify the FRT-Kan^R^-FRT cassette in pKD13. Kan^R^ cassette was amplified via PCR, DpnI-treated, and purified before electroporation into *S*. Tm IR715 harboring the helper plasmid pKD46 grown in the presence of 0.1M L-arabinose. The transformation was plated on LB agar supplemented with Kanamycin and incubated overnight at 37°C; colonies were checked by PCR. Insertion into the gene locus was confirmed using a Forward primer specific to the gene locus (i.e., *aspC*_conF) and a reverse primer specific to the KanR cassette (K1). Following insertion confirmation, the insertion was pushed into a clean background via P22 transduction as previously described^35^, then transformed with pCP20 encoding FLP to remove the Kan^R^ cassette. Clean deletion mutants were cultured at 37 °C to remove pCP20, checked for Carbenicillin sensitivity, and were PCR-verified using external primers specific to the gene locus, with WT and Kan^R^-insertion mutants as controls.

To complement the *S*. Tm ΔaspC deletion mutant, the *aspC* promoter in the 14028S genome (IR715 parental strain) was predicted using ProPr V.2 software (http://ppp.molgenrug.nl/), and primers were designed to amplify 282 bp upstream and 92 bp downstream of the *aspC* ORF via PCR. pWSK29 was linearized by inverse PCR using pWSK29_fwd and pWSK29_rev. PCR products were run on an agarose gel and purified using the Monarch PCR & DNA Cleanup Kit according to the manufacturer’s instructions. The construct was assembled using NEB HiFi assembly, transformed into electrocompetent DH5α *E. coli*, and colonies were recovered on LB+Carb. Constructs were confirmed for insertion by PCR, sequence-verified, and then transformed into the Δ*aspC S*. Tm clean deletion mutant; colonies were recovered on LB+Carb. For a pEV control, WT and Δ*aspC S*. Tm were transformed with empty pWSK29, and colonies were recovered on LB+Carb.

#### Growth Kinetics

Bacteria were streaked out onto fresh selective agar plates and incubated overnight. Single colonies were taken and cultured in LB broth supplemented with appropriate antibiotics in biological triplicate for three independent experiments. Overnight cultures were then back diluted 1:100 for 1-2 hours in 5 mL of fresh LB broth supplemented with antibiotics until the bacteria reached the mid-log phase. Cultures were then washed and normalized to an OD_600_=1. Cultures were then inoculated into a final volume of 200uL per well of pre-reduced media to a final OD_600_=0.01, shaking continuously, incubated at 37, and OD_600_ was measured every hour.

#### Gene expression analysis

Bacteria were streaked out fresh onto selective agar plates and incubated overnight. Single colonies were taken, and cultures for biological replicates of n=5 in three independent experiments were grown overnight. Overnight cultures were diluted 1:100 for 2-3 hours to reach mid-log phase, harvested, and washed. Cultures were normalized and inoculated into NCE minimal media containing ±10 mM aspartate and ±40 mM nitrate, pH normalized to 7.2. Bacteria were then incubated anaerobically for 4 hours at 37°C before being harvested and RNA extracted using Max Bacterial-RNA Extraction Reagent and TRIzol, combined with the Invitrogen PureLink RNA mini-kit, following the manufacturer’s instructions.

#### In vitro competitive infections

Bacterial strains were struck out onto selective agar plates and incubated overnight. Single colonies were taken, and cultures for biological replicates (n=5) in three independent experiments were grown overnight. Overnight cultures were then diluted 1:100 for 2-3 hours to reach the mid-log phase of growth, harvested, and washed. Cultures were then normalized and inoculated into pre-reduced NCE or NCE+0.5% Porcine Gastric Mucin (type II) at a final bacterial concentration of 1 × 10^4^ CFU/mL. Samples collected before and after 18hrs anaerobic incubation at 37°C. Samples were serially diluted and plated on LB + Kan and LB + Cm; CFU were enumerated, and CI was calculated for both time points, then normalized to determine the competition index of the inoculum.

#### Mouse models of *S*. Tm infection

All animal experiments in this study were approved by the Institutional Animal Care and Use Committee at Vanderbilt University Medical Center. Female CBA were obtained from the Jackson Laboratory. The age at the beginning of the experiment was 6-7 weeks for all CBA. Conventional mice were housed in individually ventilated cages with ad libitum access to water and chow (Envigo Global 16% Protein Rodent Diet). Unless stated otherwise, a minimum of 6 mice were used based on the variability observed in previous experiments. All mice were monitored twice daily, and cage bedding was changed every 2 weeks. At the end of the experiments, mice were humanely euthanized using carbon dioxide inhalation and cervical dislocation. Animals that had to be euthanized for humane reasons before reaching the predetermined time point were excluded from the analysis.

### Intraperitoneal infection

7-week-old CBA/J mice were purchased from Jackson Labs Inc., infected with 10^4^ C.F.U. of each strain, and carried for 96 hours post-infection before sacrifice. Mice were weighed every day post-infection. At sacrifice, peritoneal lavage, liver, spleen, and intestinal contents were collected and plated for bacterial burden via serial dilution onto LB agar supplemented with either Chloramphenicol or Kanamycin, depending on the strain background.

### Intragastric infection

7-week-old CBA/J mice were purchased from Jackson Labs Inc. and infected intragastrically with 10^9^ C.F.U. of each strain for single-strain infections or with a 1:1 ratio of WT and respective isogenic mutant strains for competitive infections. Post-infection mice were maintained for ten days, weighing daily and collecting feces every other day for bacterial burden enumeration. At sacrifice, mice were euthanized, and liver, spleen, and intestinal tissue and content were taken for bacterial enumeration via homogenization and serial dilution. Tissue was taken for histopathology evaluation.

### Quantification and Statistical analysis

Statistical data analysis was performed using GraphPad PRISM. Fold changes of ratios (bacterial competitive index and mRNA levels), and bacterial numbers were transformed logarithmically prior to statistical analysis. A Student’s t-test, paired or unpaired, was used on the transformed data to determine whether differences between groups were statistically significant (p < 0.05). When more than two treatments were used, statistically significant differences between groups were determined by one-way ANOVA followed by Tukey’s HSD test (between > 2 groups).

## Acknowledgments

N.G.S. was supported by NIH T32 Training Grant (T32ES007028-46) and HHMI Gilliam Fellowship (GT15104). J.O. was supported by NIH T32 Training Grant (T32AI095202-15). K.M.J. was supported by NIH T32 Training Grant (T32GM137793-05). M.X.B is a HHMI Freeman Hrabowski Scholar. Work in M.X.B.’s laboratory was supported by VDDRC Pilot and Feasibility grant P30 058404, the NIH (R01DK131104-01 and 1R01AI168302-01A1), The Pew Charitable Trusts (2022-A-19568), and the Burroughs Wellcome Fund (022792). Figures and images were generated using BioRender Inc.

## Author contributions

Conceptualization, N.G.S., and M.X.B.; Methodology, N.G.S, and M.X.B.; Investigation, N.G.S., M.X.B., H.F.A., J.O., K.M.J.; Writing – Original Draft, N.G.S.; Writing – Review & Editing, N.G.S., M.X.B.; Funding Acquisition, M.X.B.; Resources, M.X.B.; Supervision, M.X.B.

