## Supplementary materials for "Aspartate aminotransferase is required for *Salmonella* expansion in the inflamed gut via TCA anaplerosis"

**Supplementary Table 1: Bacterial strains and plasmids used in this study.**

| **Bacterial Strain** | **Source** | **Identification** |
|---|---|---|
| *S.* Typhimurium, IR715 ATCC 14028S Nal$^R$ | Stojiljkovic et al.[47] | IR715 |
| *S.* Typhimurium, IR715 Δ*phoN*::Tn*10d*-Cm$^R$ | Faber et al.[48] | FF176 |
| *S.* Typhimurium, IR715 Δ*phoN*::Km$^R$ | Kingsley et al.[49] | AJB715 |
| *S*. Typhimurium, IR715 Δ*aspC*::Km$^R$ | This study | NS42 |
| *S*. Typhimurium, IR715 Δ*aspC* | This study | NS43 |
| *S.* Typhimurium, IR715 Δ*frdABCD* | This study | WJ28 |
| *S.* Typhimurium, IR715 Δ*frdABCD*Δ*phoN*::Tn*10d*-Cm$^R$ | This study | WJ30 |
| *S.* Typhimurium, IR715 Δ*invA*Δ*spiB* | Rivera-Chávez et al.[50] | SPN487 |
| *S.* Typhimurium, IR715 Δ*invA*Δ*spiB*Δ*aspC*::Km$^R$ | This study | NS301 |
| *S.* Typhimurium, IR715 Δ*invA*Δ*spiB*Δ*phoN*::Tn*10d*-Cm$^R$ | This study | WJ52 |
| *S.* Typhimurium, IR715 Δ*invA*Δ*spiB*Δ*aspC* | This study | NS311 |
| *S.* Typhimurium, IR715 Δ*invA*Δ*spiB*Δ*aspC*Δ*phoN::*Km$^R$ | This study | NS313 |
| *S*. Typhimurium, IR715 Δ*sdhA*::Km$^R$ | This study | NS314 |
| *S*. Typhimurium, IR715 Δ*gltA*::Km$^R$ | This study | NS315 |
| *S*. Typhimurium, IR715 Δ*sdhA*Δ*aspC*::Km$^R$ | This study | NS316 |
| *S*. Typhimurium, IR715 Δ*gltA*Δ*aspC*::Km$^R$ | This study | NS317 |
| *S*. Typhimurium, IR715 Δ*sdhA*ΔphoN::Tn10d-Cm$^R$ | This study | NS318 |
| *S*. Typhimurium, IR715 Δ*gltA*Δ*phoN*::Tn10d-Cm$^R$ | This study | NS319 |
| *S*. Typhimurium, IR715 Δ*sdhA*Δ*aspC*Δ*phoN*::Km$^R$ | This study | NS320 |
| *S*. Typhimurium, IR715 Δ*gltA*Δ*aspC*Δ*phoN*::Km$^R$ | This study | NS321 |
| *S*. Typhimurium, IR715 WT:pWSK29 (pEV) | This study | HA109 |
| *S*. Typhimurium, IR715 Δ*aspC*:pWSK29 (pEV) | This study | HA113 |
| *S*. Typhimurium, IR715 Δ*aspC*:pWSK29::*aspC* | This study | HA114 |
| **Plasmid** | **Source** | **Identification** |
| Plasmid: pKD46, Spec$^R$ $P_{BAD}$-*gam-beta-exo oriR101 repA101*$^{ts}$ | Datsenko and Wanner[54] | pKD46 |
| Plasmid: pKD3, Carb$^R$ FRT Cm$^R$ FRT PS1 PS2 *oriR6Kγ* | Datsenko and Wanne[54] | pKD3 |
| Plasmid: pKD13, Carb$^R$ FRT Km$^R$ FRT PS1 PS4 *oriR6Kγ* | Datsenko and Wanner[54] | pKD13 |
| Plasmid: pCP20, CarbR FLP *oripSC101* CmR r*epA101*$^{ts}$ MbeC | Datsenko and Wanner[54] | pCP20 |
| Plasmid: pWSK29, CarbR LacZα *rep101* pSC101ori | Murray GL[55] | pEV |

| | | |
|---|---|---|
| Plasmid: pWSK29::*aspC,* CarbR LacZ$\alpha$ *rep101* pSC101ori *aspC* | This study | pWSK29::*aspC* |

**Supplementary Table 2: Primers used in this study.**

| Primer | Sequence |
|---|---|
| IR715_gltA_delF | GCTATTGAACTGGATGTGCTAAAAGGTACACTCGGTCAATGTAGGCT GGAGCTGCTTCG |
| IR715_gltA_delR | GATTTTCATGCCGTCAGTGTGCATTTCATTCCAGTGTGCATTCC GGG GATCCGTCGACC |
| IR715_gltA_conF | GGTCACTCTTTCACCTGTTA |
| IR715_gltA_conR | GCCGGAAGGTGCAGATAGAG |
| IR715-sdhA-del-F1 | CAAACCTGTGCGCTGCTCTCGAAAGTCTTCCCGACCCGTTTGTAGG CT GGAGCTGCTTCG |
| IR715-sdhA-del-R1 | CTGATACAGGGTATGACACACCA GTTGGCATCATCACGCATTCCGGGGATCCGTCGACC |
| IR715-sdhA-con-F1 | GCAACGCCTCCGCATTAGGA |
| IR715-sdhA-con-R1 | CACACCTTCGCGGCAGGAAC |
| aspC-del-F1 | ATAGCGGATTTCCCTTCTGTAACTATAATGGACCT CGTCTGTAGGCT GGAGCTGCTTCG |
| aspC-del-R1 | ATTGCAGGCTTTTTTATCGGTCACGCCAGTCGGCAGCTTTATTCCGG G GATCCGTCGACC |
| aspC-con-F1 | TACTTACCGTT CTGATCAGC |
| aspC-con-R1 | CGCTTGGCGATGATGTGAC |
| aspC_comp_Fwd | ATCAAGCTTATCGATACCGTC |
| aspC_comp_Rev | ATCGAATTCCTGCAGCCC |
| pWSK29_Fwd | CCGGGCTGCAGGAATTCGATTGACTACCGTTTCAACCTG |
| pWSK29_rev | ACGGTATCGATAAGCTTGATCTTCTTGCAAGATGAGATG |
| thrA_qFwd | GGCGTTCTACAGCCACTATTATC |
| thrA_qRev | CCGTAACAGATCGGCAAACA |
| metL_qFwd | GGGCGTTACTGGCTGTATAA |
| melL_qRev | CGCTGATCGAGAGAATCGTATC |
| lysC_qFwd | GGTACGCTGGTGTGCAATAA |
| lysC_qREv | CCTCGTGAGTGCAGCATATT |
| pyrI_qFwd | AACCAGTATCCTCCAGCTTTG |
| pyrI_qRev | TTGGCCAGCACCACATAAT |
| pyrB_qFwd | CAGCCGCGACGATCTTAAT |
| pyrB_qRev | GCTGGCGATCACTTTATGTTTC |
| asnA_qFwd | GAGGCTATATGGGCAGGAATTA |
| asnA_qRev | CAGTTCCTGACTGTGAACAAAC |
| IR715_qPCR_aspA_F | TCGTAACGGTGGTGTCGTTA |
| IR715_qPCR_aspA_R | CTGTGGCGCATATGTTATGG |
| IR715_qPCR_aspC_F | CAGACCGCTGAATGAGAACA |
| IR715_qPCR_aspC_R | GCCATTCGCGCTAACTACTC |
| IR715_qPCR_dcuA_F | GATGACCCTGGATGACGCTA |
| IR715_qPCR_dcuA_R | AAACCGCTGATGAACACCAC |
| K1 | CAGTCATAGCCGAATAGCCT |

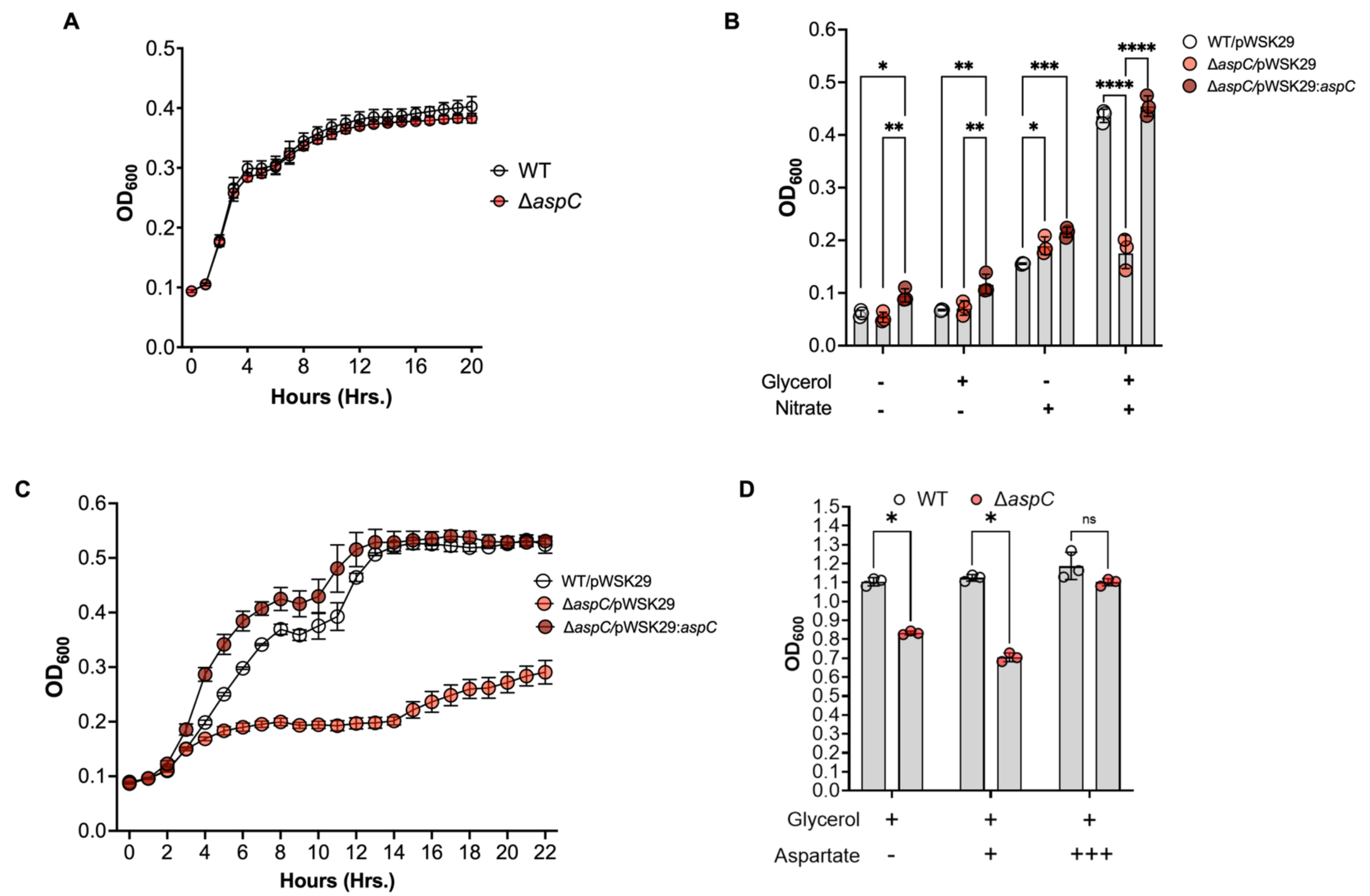


**Figure S1: Deletion of Δ*aspC* does not confer a growth defect compared to WT when cultured anaerobically in LB broth but does under hypoxia in minimal media.** Related to Figure 4. (A) Growth of WT and *aspC*-deficient *S*. Tm (Δ*aspC*) anaerobically in LB. (B) Growth kinetics of WT (WT/pWSK29), *aspC*-deficient (Δ*aspC*/PWSK29), and complemented *aspC*-deficient *S.* Tm (Δ*aspC*/pWSK29*:aspC*) under anaerobic conditions in NCE media supplemented with glycerol (40 mM) and/or nitrate (40 mM) were measured at 18 hours post-inoculation. (C) Growth kinetics of WT (WT/pWSK29), *aspC*-deficient (Δ*aspC*/PWSK29) and complemented *aspC*-deficient *S.* Tm (Δ*aspC*/pWSK29*:aspC*) under anaerobic conditions in NCE media supplemented with glycerol (40 mM) and nitrate (40 mM). (D) Growth of WT and *aspC*-deficient *S*. Tm ((Δ*aspC*) under hypoxic (0.8% $O_2$) NCE minimal media supplemented with Glycerol and (+) 0.1mM or (+++) 10mM Aspartate with Glycerol. N=3 per condition. Geometric mean and Geometric SD. *, p<0.05; **, p<0.01; ***, p<0.001 using 2-way ANOVA (B) and multiple unpaired t-test (D).

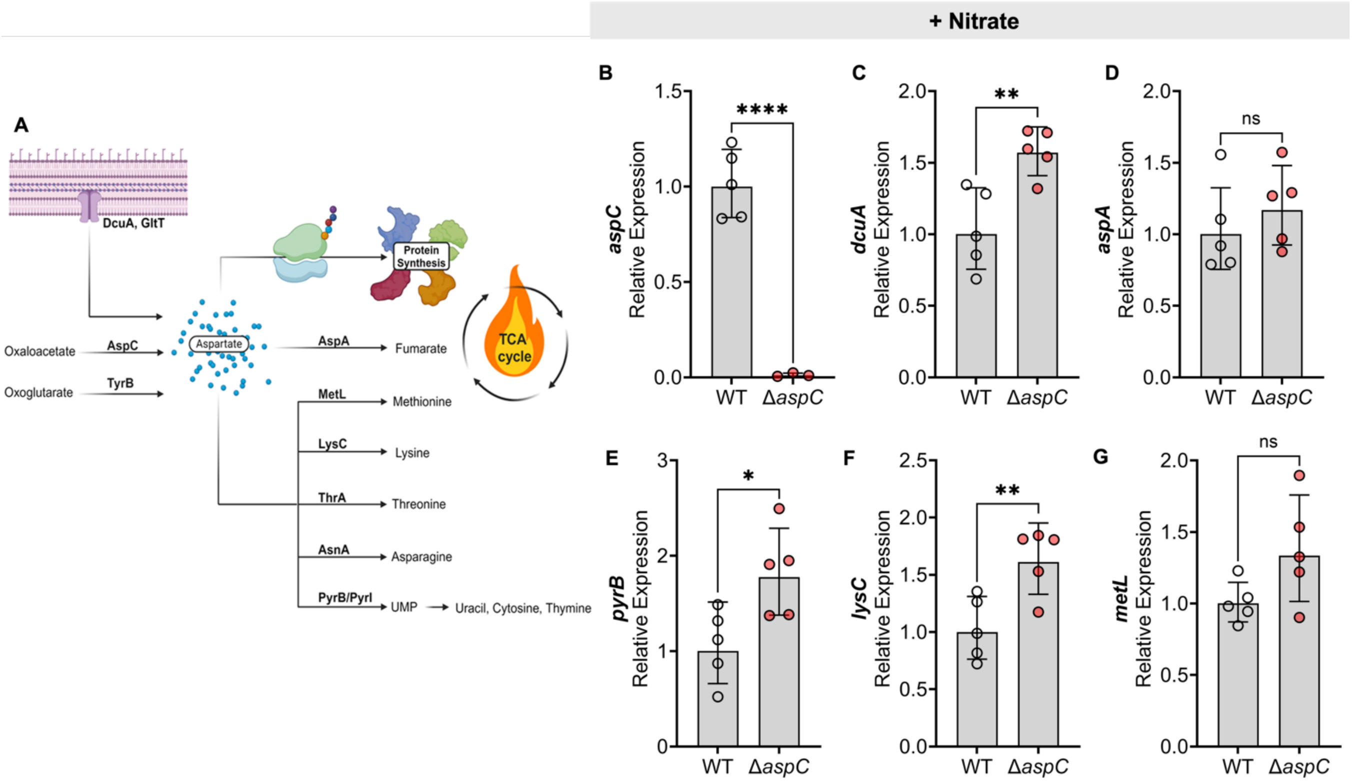


**Figure S2: AspC-deficiency alter central metabolism of *S.* Tm during anaerobic nitrate respiration.** Related to Figure 5. As aspartate can fuel several other metabolic processes, we investigated the expression of genes involved in aspartate utilization in conditions supplemented with nitrate, in which we observed a significant growth and fitness defect of the *aspC*-deficient *S.* Tm (Δ*aspC*) compared to the WT. (A) Schematic for the central role of aspartate in *S.* Tm metabolism. (B-G) Gene expression of WT and Δ*aspC* grown anaerobically under nitrate respiration (40 mM nitrate) with glycerol (40 mM) as the sole-carbon source. (B) *aspC,* (C) *dcuA,* (D) *aspA,* (E) *pyrB*, (F) *lysC*, (G) *metL*. N=5 per condition, Geometric mean and Geometric SD. *, p<0.05; **, p<0.01; ***, p<0.001; ****, p<0.0001 using unpaired t-test.

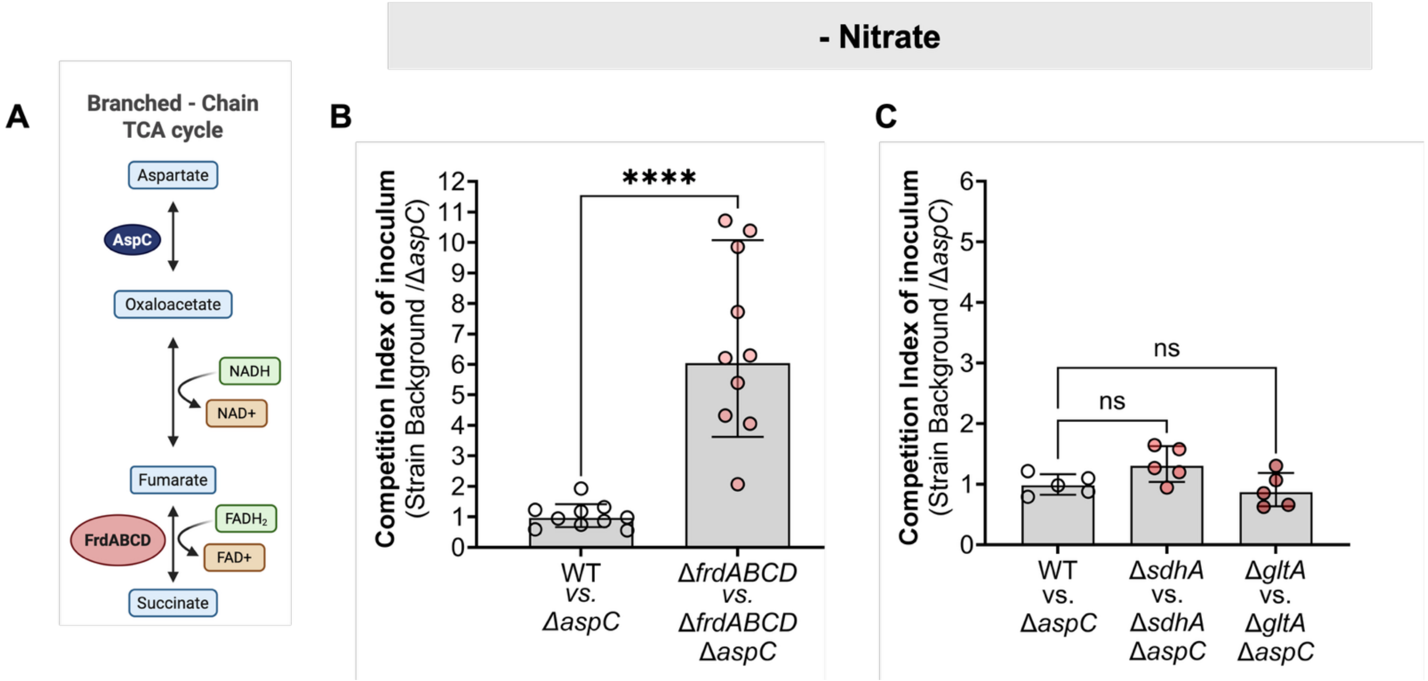


**Figure S3: Deletion of *frdABCD* but not *sdhA* or *gltA* significantly increases the fitness defect of *aspC*-deficient *S.* Tm during *in vitro* growth in mucin media alone.** Related to Figure 5. (A) Competitive index of inoculum for loss of *aspC (ΔaspC)* in WT and *frdABCD*–deficient *(ΔfrdABCD) S*. Tm backgrounds anaerobically grown in mucin media for 18 hours in the absence of nitrate**.** (B) Competitive index of inoculum for loss of *aspC (ΔaspC)* in WT and *gltA (ΔgltA) or sdhA (ΔsdhA)*–deficient backgrounds anaerobically grown in mucin media for 18 hours in the absence of nitrate**.** N=5-10 per condition. Geometric mean and Geometric SD. ****, $p<0.0001$ using unpaired t-test.
